# Microparasite prevalence in Southern giant pouched rats (*Cricetomys ansorgei*) in Morogoro, Tanzania

**DOI:** 10.1101/2025.07.24.666516

**Authors:** Cato Vangenechten, Joachim Mariën, Hélène Vandecasteele, Natalie Van Houtte, Baraka E. Mwamundela, Christopher Sabuni, Sophie Gryseels, Herwig Leirs, Lucinda Kirkpatrick

## Abstract

Rodents play a critical role in the transmission of zoonotic pathogens, with environmental changes and increasing human-wildlife interactions further amplifying disease spillover risks. *Cricetomys* spp., commonly found in both human dwellings and agricultural fields, are frequently hunted for consumption, potentially facilitating pathogen transmission. However, their roles in pathogen transmission and disease ecology remain poorly understood. This study investigated the prevalence of three microparasite genera, i.e., *Bartonella, Hepatozoon*, and *Anaplasma*, in *Cricetomys* spp. from Morogoro, Tanzania. We observed a prevalence of 71.9% for *Bartonella*, 17.5% for *Hepatozoon*, and 1.8% for *Anaplasma*. To our knowledge, this is the first study to investigate *Bartonella, Hepatozoon*, and *Anaplasma* in *Cricetomys* spp. in Tanzania. Furthermore, preliminary results indicate that *Bartonella* infection prevalence is influenced by habitat type, with a significantly higher prevalence observed in rural areas compared to urban areas. This study underscores the potential role of *Cricetomys* spp. as important reservoirs of infectious diseases.

## 1 Introduction

Over the past five decades, there has been a notable increase in the number of emerging infectious diseases (EIDs) worldwide, driven by factors such as climate change, shifts in human demographics, and increased globalization (Jones et al. 2008; van Doorn 2017). These factors can directly or indirectly affect disease dynamics, for instance by causing biodiversity declines through the overexploitation of native species and habitat degradation (Pimm and Raven 2000), or by increasing human-animal contact through human encroachment into previously undisturbed areas (Gibb et al. 2020; Mangombi et al. 2021; Newbold et al. 2015; Panthawong et al. 2020). Furthermore, deteriorating environmental conditions may elevate stress on wildlife, increasing their vulnerability to infections by zoonotic microparasites, and thereby heightening the risk of transmission to humans (Ostfeld and Keesing 2012; Reaser et al. 2021).

Rodents represent the largest order of mammals, comprising approximately 42% of the global mammalian biodiversity, and are among the few animal groups that thrive in close association with humans (Bordes et al. 2015; Dahmana et al. 2020). In the context of ongoing environmental changes, many rodent species are expanding beyond their natural distribution ranges, largely due to their synanthropic tendencies (Dahmana et al. 2020). Certain biological characteristics of these rodents, like early maturation, large litter sizes, short lifespans, and the prioritization of reproduction over immunity, make them common reservoirs for zoonotic pathogens (Chicago 2007; McCauley et al. 2015; Suzán et al. 2009). Indeed, rodents are known to host over 60 zoonotic diseases, including plague, Lassa fever, and hantavirus-related diseases, many of which occur in the Afrotropics (Dahmana et al. 2020). Human encroachment into previously undisturbed areas increases the likelihood of direct contact with wildlife or indirect contact through contaminated faeces or urine (Gibb et al. 2020; Mangombi et al. 2021; Newbold et al. 2015; Panthawong et al. 2020). Furthermore, the hunting and consumption of rodents presents significant risks for cross-species transmission of zoonotic pathogens (Wolfe et al. 2005). To date, the potential for disease emergence through wildlife consumption remains a global concern due to increasing human population density, globalized trade, and heightened human-animal interactions. In many rural regions of West and Central Africa, bushmeat is an important food source. In the urban areas and diaspora communities, bushmeat delicacies are also in high demand, increasing the hunting pressure beyond sustainability (Fa and Brown 2009; Ingram et al. 2025; Lucas et al. 2022). This hunting, trade, and consumption of wildlife, including rodents, elevates human exposure to zoonotic pathogens, increasing the likelihood of disease spillover events (Milbank and Vira 2022; Wolfe et al. 2005).

*Cricetomys* spp., commonly known as African giant pouched rats, are among the rodent genera frequently consumed as bushmeat. This study focuses on microparasite infections in *Cricetomys ansorgei*, a rodent species found throughout tropical and subtropical regions in sub-Saharan Africa (Katakweba et al. 2022). These rats predominantly inhabit terrestrial environments, constructing underground burrows that are observed not only in dense forests, but also in farmlands, fallow fields, and near abandoned houses (Kalemba et al. 2022). *Cricetomys* spp. are also commonly encountered in urban areas, where they live in close proximity to humans (Katakweba et al. 2022). They are solitary and territorial, with burrows typically occupied by a single individual, though females with juveniles may cohabitate for extended periods (Kalemba et al. 2022). Their broad ecological distribution – across urban, agricultural, and forested environments – combined with their value as a food source, underscores their relevance in zoonotic disease ecology (Morand et al. 2019; van Vliet et al. 2012). Previous studies indicate that *Cricetomys* spp. may serve as hosts for several zoonotic pathogens, including *Leptospira, Bartonella, Trypanosoma, Mycobacteria, Toxoplasma*, and orthopoxviruses (Billeter et al. 2014; Demoncheaux et al. 2022; Durnez et al. 2008; Galal et al. 2019; Mangombi et al. 2021; Mgode et al. 2015; Reynolds et al. 2010). To date, *Cricetomys* spp. remain understudied, partly due to the difficulties associated with their capture and the need for specialized equipment and expertise. These challenges, combined with limited research funding and prioritization, contribute to the scarcity of data on these rodent species.

This study investigates the prevalence of three microparasite genera, i.e., *Bartonella, Anaplasma*, and *Hepatozoon*, as they are commonly found in rodents across sub-Saharan Africa. Moreover, these pathogens have significant implications for both human and animal health. (i) *Bartonella* is a genus of small, gram-negative bacteria responsible for causing human diseases such as trench fever (*Bartonella quintana*), cat scratch disease (*Bartonella henselae*), and Carrion’s disease (*Bartonella bacilliformis*) (Gutierrez et al. 2015). *Bartonella* primarily targets mammalian erythrocytes and endothelial cells (Theonest et al. 2019), and is transmitted by hematophagous arthropods such as fleas, lice, and ticks (Billeter et al. 2008). (ii) *Anaplasma* spp. are gram-negative, obligate intracellular bacteria that infect mammalian blood cells (Rymaszewska and Grenda 2008). Species of this genus may infect humans and various animals, causing conditions like human granulocytic anaplasmosis (*A. phagocytophilum*), bovine anaplasmosis (*A. marginale*), monocytic anaplasmosis (*A. bovis*), anaplasmosis in small ruminants (*A. ovis*), and canine cyclic thrombocytopenia (*A. platys*) (Rymaszewska and Grenda 2008). The life cycle of *Anaplasma* spp. involves both vertebrate and invertebrate hosts, with ticks serving as the primary vectors (Silaghi et al. 2017). (iii) *Hepatozoon* comprises apicomplexan protozoa that parasitize a diverse range of hosts, including birds, mammals, lizards, and snakes (Smith 1996). Their heteroxenous life cycle involves vertebrate intermediate hosts and hematophagous invertebrates as definitive hosts. In domestic animals, *Hepatozoon* infections often result in hepatozoonosis (Baneth and Allen 2022).

The aim of this study is to assess the prevalence of the selected microparasites in *Cricetomys* spp. to better understand the potential role of these rodents as reservoirs for zoonotic pathogens. Given the close association between rodents and human populations, understanding their capacity to harbour these microparasites is critical for evaluating public health risks. By addressing this knowledge gap, this study contributes to a broader understanding of disease transmission dynamics and the potential spillover risk associated with human and rodent interactions.

## 2 Material and methods

### 2.1 Study site and trapping

*Cricetomys* spp. were trapped at fourteen sites in Morogoro, Tanzania, during July and August 2022 (Figure 1). Selection of the study sites was conducted prior to the start of the research based on specific criteria: (i) the presence of water bodies, large rocks, and trees; and (ii) evidence of *Cricetomys* activity, such as faeces, along with confirmation from local inhabitants regarding the presence of *Cricetomys*. The capture sites were categorized into two groups, i.e., urban and rural areas, based on land cover data which was derived from an existing land classification database (ESA World Cover; 10m; 2021; v200) and ground truthed before traps were set. Havahart® Live Animal Traps (Havahart®, Pennsylvania, USA), baited with a mixture of banana maize flour and roasted sardine fish, were strategically positioned in the late afternoon and checked the following morning. Captured *Cricetomys* were brought to the Institute of Pest Management Centre (IPM) at the Sokoine University of Agriculture (SUA). Geographic coordinates were documented for each trapping location (Supplementary Table S1). Upon capture, the *Cricetomys* were individually housed in cages with bedding material. Euthanasia and dissections were conducted promptly after capture to minimize stress and to ensure that infections could not occur post-trapping.

**Figure 1.**
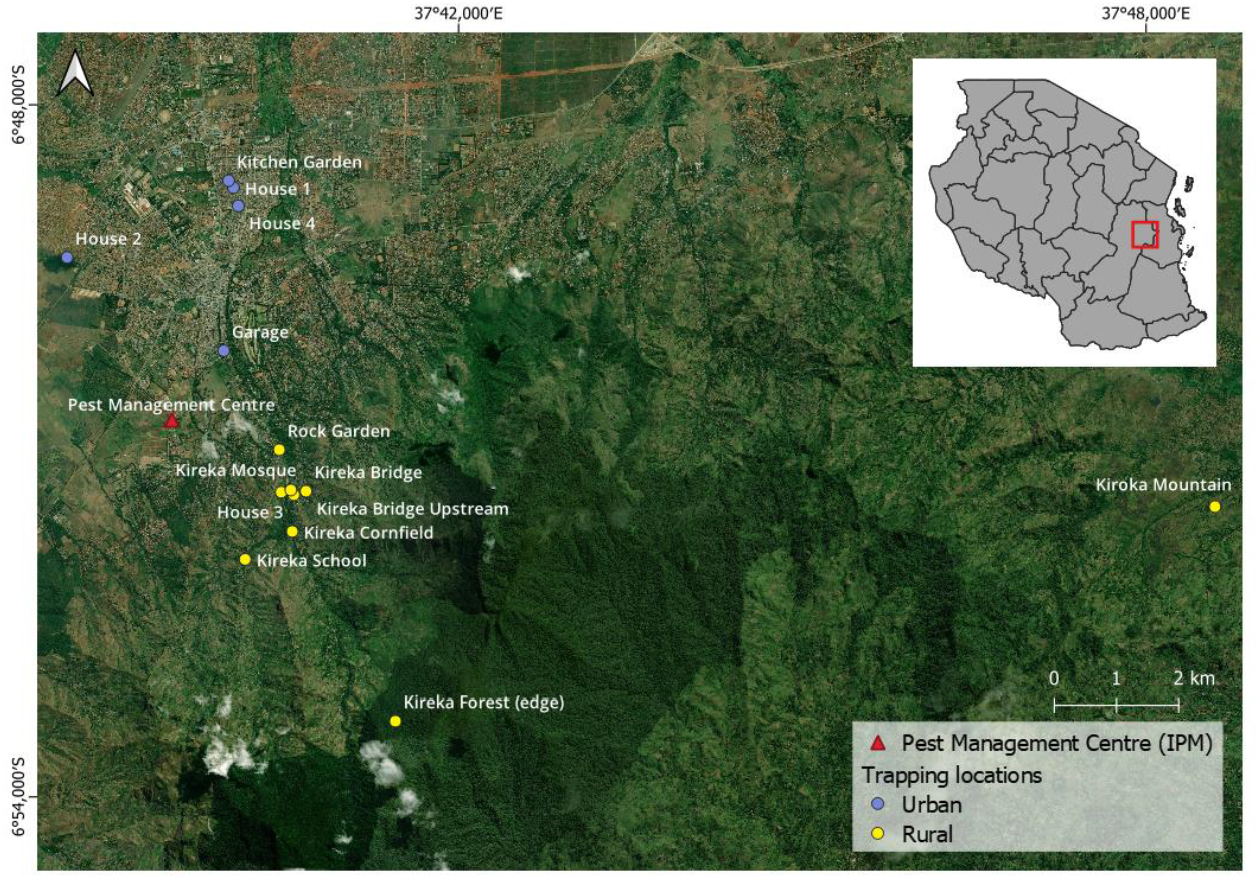
Map of Morogoro, Tanzania, illustrating the trapping locations of *Cricetomys* spp. and the Institute of Pest Management Centre (IPM). Coloured circles indicate the different habitat types of the trapping locations (urban = blue and rural = yellow). The inset map shows the outline of Tanzania, with the study site marked by a red square for reference within the country.

All individuals underwent comprehensive biometric assessments, including recording of body weight, sex, reproductive status, body length (cm), and tail length (cm). Euthanasia was performed using an overdose of isoflurane followed by cervical dislocation. Blood samples were collected directly from the heart and deposited onto Whatman filter paper (Sigma-Aldrich, Missouri, USA) for preservation. Spleen tissue samples were dissected and stored in DNA/RNA shield for preservation, and eyes were preserved in 10% formalin. To avoid cross-contamination, scissors and tweezers were sterilized by sequentially immersing them in 10% bleach, rinsing with distilled water, immersion in 100% ethanol, and flame sterilization. The age of the captured *Cricetomys* spp. was estimated based on the eye lens weight (ELW), which increases with age independently of external factors (Morris 1972). Eye lenses were extracted from the eyes using tweezers, rinsed with distilled water, and dried for two hours at 100°C. After drying, the eye lenses were weighed to the nearest 0.001 g (Stephenson et al. 1984).

### 2.2 Molecular detection of pathogenic DNA

DNA was extracted from the spleen using the Quick-DNA/RNA^TM^ Pathogen Miniprep kit (Zymo Research, Irvine, USA) following the manufacturer’s protocol. To prevent contamination when handling tissues from different animals, all instruments, including scalpels, scissors, and tweezers, were decontaminated with 5% Virkon® (Antec International, Sudbury, UK) and allowed to dry after processing each tissue sample. The quality and quantity of DNA extracts were assessed using the Qubit fluorometer (Thermo Fisher Scientific, Massachusetts, USA). Subsequently, all DNA extracts were stored at −20 °C until PCR analysis.

Firstly, DNA extracts were screened on the presence of *Bartonella* and *Anaplasma* spp. using the StepOne^TM^ Real-Time PCR system (Applied Biosystems, Massachusetts, USA) (Table 1) (Dahmana et al. 2020). Amplification reactions were conducted in a final volume of 15 µL containing 7.50 µL of Takyon^TM^ ROX 2X MasterMix dTTP (Eurogentec, Liège, Belgium), 0.75 µL of the forward and reverse primer (final concentration 0.50 µM), 0.36 µL of the probe (final concentration 0.12 µM), 3.14 µL of DNase-free water, and 2.50 µL of DNA template. The following thermal profile was used: incubation at 50°C for two minutes, an activation step at 95°C for three minutes, followed by 40 denaturation cycles at 95°C for 15 seconds and an annealing-extension at the annealing temperature of the corresponding primers for 30 seconds. A positive and negative control were added to each PCR run. Samples with a CT-value below 35 were screened a second time (duplicates). A sample was considered positive in the qPCR if it tested positive in both qPCR runs.

**Table 1.**
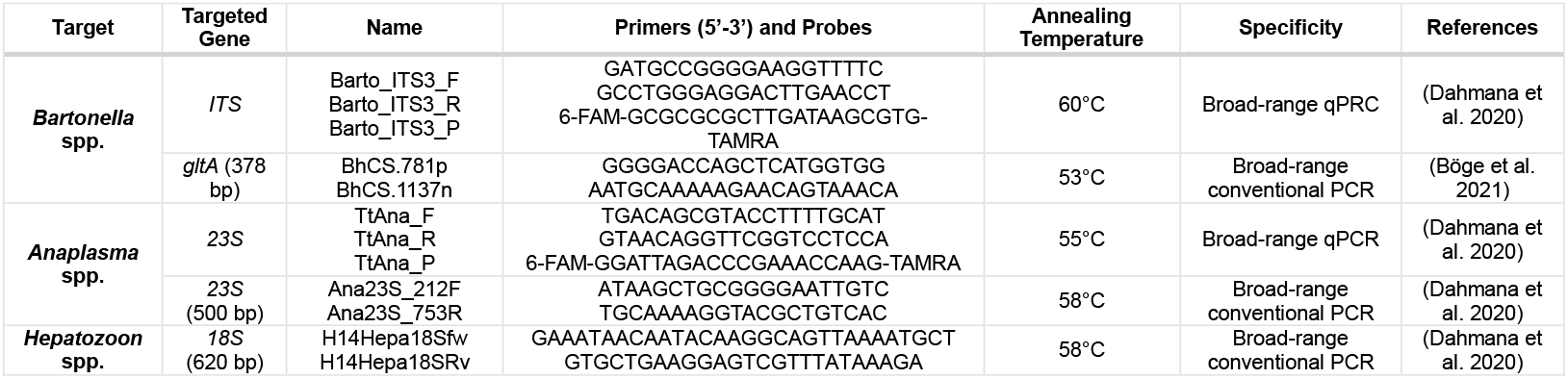
Oligonucleotide sequences of primers and probes used for qPCRs and conventional PCRs.

Secondly, conventional PCR was used to amplify all qPCR positive samples for *Bartonella* spp. targeting the *gltA* gene, for *Anaplasma* spp. targeting the *23S* gene, and for *Hepatozoon* spp. targeting the *18S* gene (Table 1) (Böge et al. 2021; Dahmana et al. 2020). Amplification reactions for the detection of *Anaplasma* and *Hepatozoon* spp. were conducted in a final volume of 25 µL containing 12.50 µL of 2X-Gold STAR^TM^Mix (hot start; Eurogentec, Liège, Belgium), 0.75 µL of each primer (final concentration 0.50 µM), 8.50 µL of DNase-free water, and 2.50 µL of DNA template. For *Bartonella* spp., amplification reactions consisted of 12.50 µL of 2X-Gold STAR^TM^Mix (hot start; Eurogentec, Liège, Belgium), 0.50 µL of each primer (final concentration 0.50 µM), 9 µL of DNase-free water, and 2.50 µL of DNA template. Reactions were performed in an automated thermal cycler (TProfessional Basic Thermocycler by Analytik Jena, Jena, Germany) following the specific thermal cycling profile. For *Anaplasma* and *Hepatozoon* spp.: one incubation step at 95°C for 15 minutes, 40 cycles of 60 seconds at 95°C, 30 seconds at annealing temperature and one minute at 72 °C and a final extension step of 5 minutes at 72°C. Amplification of the *gltA Bartonella* gene consisted of 45 cycles of denaturation at 95°C for 30 seconds, annealing at 53°C for 30 seconds and elongation at 72°C for one minute.

Lastly, we assessed the quality of the PCR products through gel electrophoresis (1.4% agarose gel, 100 V for 30 min). Amplicons of positive samples were purified and sequenced by the Neuromics Support Facility, located in Antwerp, Belgium, using the forward and reverse primer. Samples were considered positive if Sanger sequences were obtained. In case no sequences were obtained, the samples that were repeatedly positive in RT-qPCR were considered uncertain.

### 2.3 Phylogenetic analysis

All sequences were edited using Geneious Prime 2024.0.5. and deposited in GenBank under the accession numbers PQ142944-PQ142977 and PQ149217-PQ49218 for *Bartonella*, PP980729-PP980738 for *Hepatozoon*, and PP980726 for *Anaplasma*. We explored the genetic diversity of the sequences with BLASTn, followed by alignment of the sequences with reference sequences from GenBank. Positions containing gaps and missing data were removed using the ‘Mask Alignment’ function in Geneious Prime 2024.0.5. Next, phylogenetic trees were constructed using maximum likelihood inference with 1000 bootstrap replicates, while assessing different substitution models in IQ-TREE 2.4.0. For the *Bartonella* dataset, the Hasegawa-Kishino-Yano model (i.e. HKY+F+G4) was selected, while for *Anaplasma* and *Hepatozoon*, the Tamura-Nei method (i.e. TN+F+I) was applied (Minh et al. 2020). Low-quality sequences containing ambiguities were excluded from the phylogenetic analysis to minimize potential biases. The phylogenetic trees were visualized in iTOL version 5 (Letunic and Bork 2021).

### 2.4 Statistical analysis

To test for significant differences in strain composition between urban and rural areas, we conducted a permutational multivariate analysis of variance (PERMANOVA) using ‘adonis2()’ function (R package ‘vegan’, 999 permutations, RStudio 2023.12.1), with habitat type as the independent variable.

To examine the relationship between habitat type and pathogen prevalence in *Cricetomys* spp., we implemented a spatial generalized linear model (GLM) using the Integrated Nested Laplace Approximation (INLA) approach within a Bayesian framework to account for potential spatial autocorrelation, as samples were collected from different geographic areas. Spatial structure was incorporated through a stochastic partial differential equation (SPDE) approach (Bakka et al. 2018), which models spatially correlated random effects as a continuous Gaussian random field (GRF) over the study area. The SPDE framework represents the GRF via a triangulated mesh, allowing computationally efficient approximation of spatial covariance (Rue et al. 2009). This method captures spatially structured variation in the response variable that is not explained by the fixed effects, improving model accuracy and accounting for potential spatial confounding. Pathogen infection status was modelled using a binomial distribution, with habitat type (urban, rural), sex (male, female) and age (ELW) as explanatory variables. All models were fitted using the INLA package in R (RStudio 2023.12.1) (Rue et al. 2009).

We have excluded individuals with an uncertain microparasite status from all statistical analyses to minimize potential misclassification bias.

## 3 Results

### 3.2 PCR results

We trapped a total of 57 individuals (N_male adult_ = 30, N_female adult_ = 20, and N_juvenile_ = 7). Of these, 21 were captured in urban areas and 36 in rural areas. In total, 41 individuals tested positive for *Bartonella* (N_rural_ = 30 and N_urban_ = 11) and 10 individuals tested positive for *Hepatozoon* (N_rural_ = 8 and N_urban_ = 2) (Figure 2). Only one individual, captured in a rural area, tested positive for *Anaplasma*.

**Figure 2.**
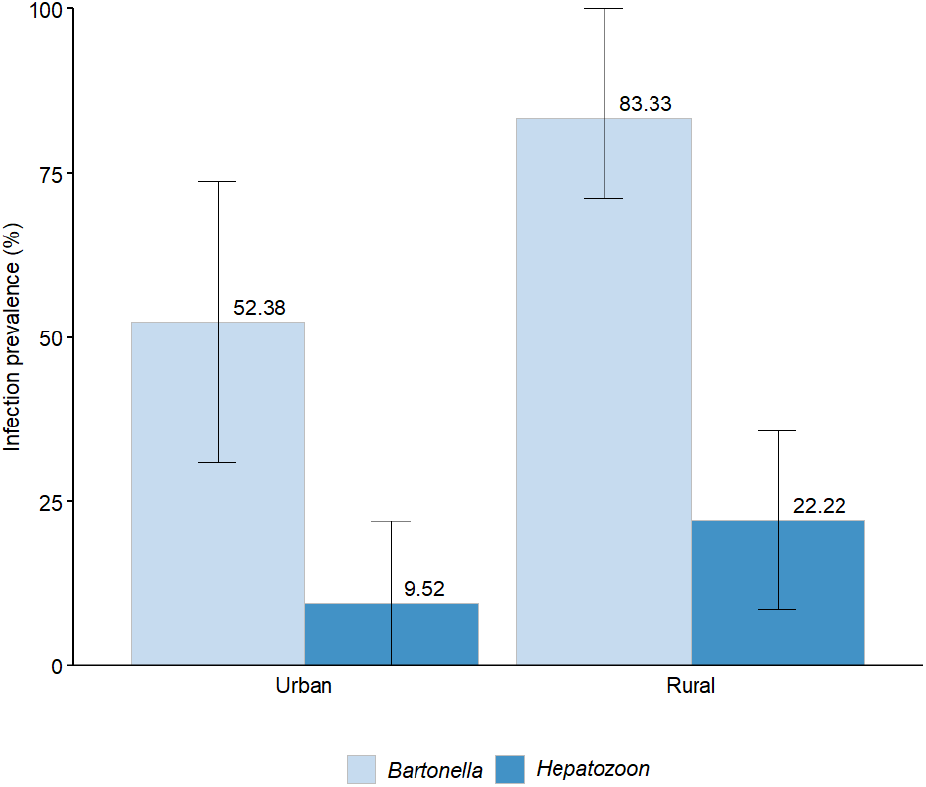
Prevalence of *Bartonella* and *Hepatozoon* infections across different habitat types, presented with 95% confidence intervals. The infection prevalence is depicted by coloured bars (*Bartonella* = light blue and *Hepatozoon* = dark blue), with the exact prevalence values annotated above each corresponding bar. The error bars represent the 95% confidence intervals.

*Bartonella* sequences were obtained for 41 individuals (71.9%, 95% CI: 60.2-83.6%; Figure 3A). For three individuals, we did not obtain a sequence despite positive results in the RT-qPCR assay; these individuals were classified as uncertain for *Bartonella* status. The phylogenetic analysis was conducted using sequences of 220 bp in length, following alignment and quality filtering. Nine sequences were excluded due to low sequencing quality to ensure the accuracy of the phylogenetic reconstruction. In total, 21 sequences showed the highest similarity in GenBank to *Bartonella* sp. SC-tr1 (AJ583132, 95.5-99.0%), previously found in *Saccostomys campestris* in South Africa (Pretorius et al. 2004). Twelve sequences were most closely related to *Bartonella* sp. Cg4ug (JX428750, 97.9-99.3%), and one sequence to *Bartonella* sp. Cg1ug (JX428749, 99.7%), both isolated from *Cricetomys gambianus* in Uganda (Billeter et al. 2014). One sequence was most similar to *Bartonella* sp. AN-tr103 (AJ583118, 97.3%), isolated from *Aethomys namaquensis* in South Africa (Pretorius et al. 2004), and another to *Bartonella massiliensis* (HM636447, 99.3%), previously found *Ornithodoros sonrai* in Senegal (Mediannikov et al. 2014).

**Figure 3.**
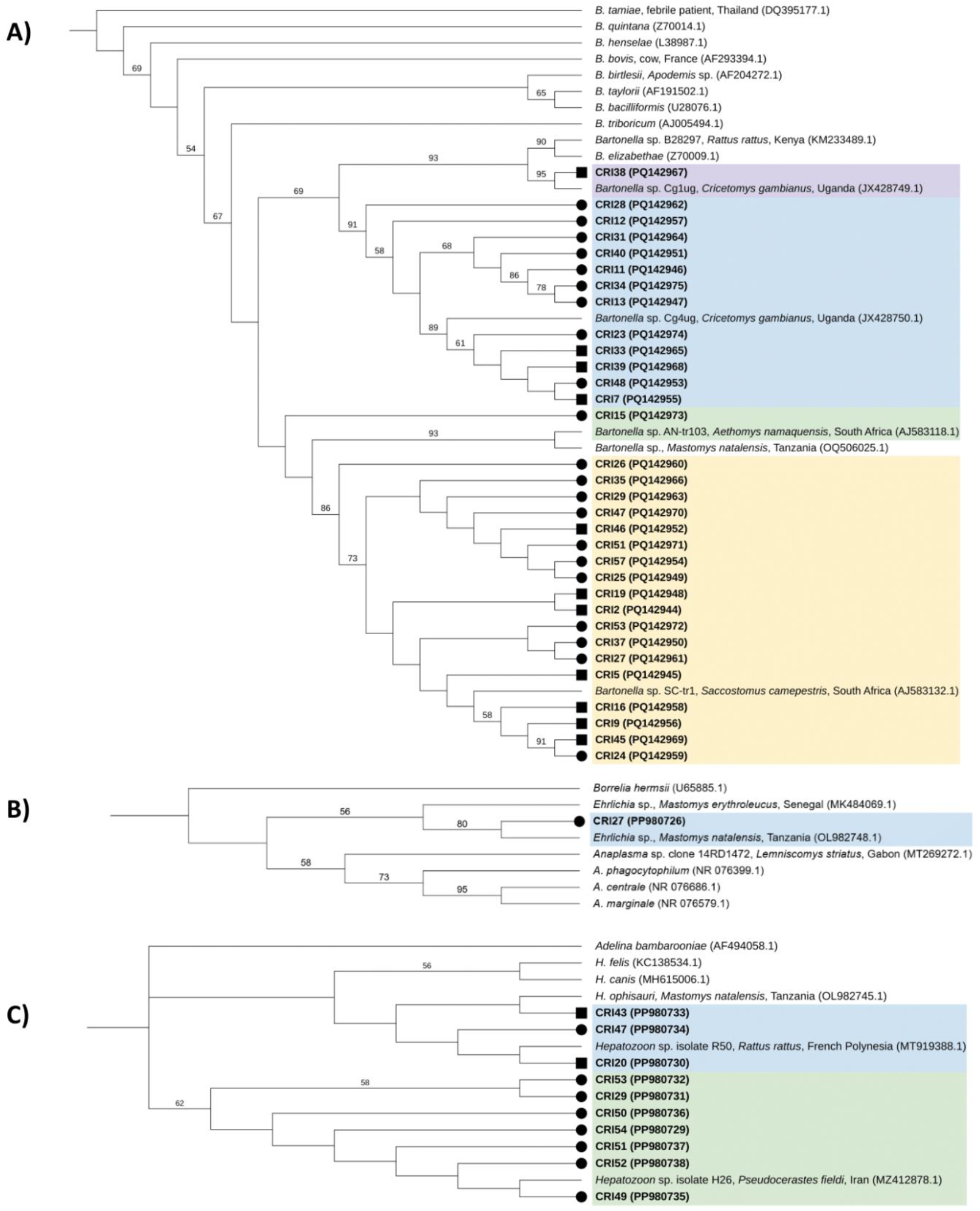
Cladogram of (A) *Bartonella gltA* sequences, (B) *Anaplasma* and *Ehrlichia* 23S rRNA sequences, and (C) *Hepatozoon* 18S sequences derived from spleen tissue samples of *Cricetomys* spp. captured in Morogoro, Tanzania. Bootstrap values greater than 50% are indicated above the corresponding branches. The sequences from this study are highlighted in bold and labelled with the prefix “CRI” followed by sample identifier numbers, and their respective GenBank accession numbers in parentheses. Capture locations are represented by symbols (urban = square and rural = circle). Reference strain sequences are included with the species name, host, and country, and GenBank accession numbers in parentheses. The sequence colours indicate groups comprising sequences obtained in this study along with their closest matches in GenBank. The outgroups used for the phylogenetic analysis were *B. tamiae* (DQ395177.1) for *Bartonella* spp., *Borrelia hermsii* for *Anaplasma* spp. (U65885.1), and *Adelina bambarooniae* (AF494058.1) for *Hepatozoon* spp. A phylogram of the same phylogenetic tree (with branch lengths proportional to phylogenetic distance) is displayed in Supplementary Figure S2.

For *Anaplasma*, one individual tested positive (1.8%, 95% CI, 0-5.2%; Figure 3B). This sequence showed the highest similarity to an uncultured *Ehrlichia* sp. (OL982748, 99.8%), originally found in *Mastomys natalensis* from the same research area (Vanden Broecke et al. 2023). The phylogenetic analysis was based on sequences of 264 bp.

In total, 10 individuals tested positive for *Hepatozoon* spp. (17.5%, 95% CI, 7.7-27.4%) (Figure 3C). Sequences were obtained for all positive individuals and phylogenetic analysis was conducted using sequences of 483 bp. Seven sequences showed the highest similarity to a *Hepatozoon* sp. isolate (MZ412878, 98.6-100%), previously isolated from *Pseudocerastes fieldi* in Iran (Jameie et al. 2022). The remaining three sequences were most similar to another *Hepatozoon* sp. isolate (MT919388, 96.9-99.5%), originally found in *Rattus rattus* in French Polynesia (Hrazdilová et al. 2021) (Supplementary Figure S2 and Supplementary Table S3).

### 3.2 Statistical analysis

The PERMANOVA analysis showed no significant differences in the composition of *Bartonella* strains between the two habitat types, i.e., urban and rural (df = 1, F = 0.99, *p* = 0.72). Furthermore, we employed a generalized linear model to examine the relationship between habitat type and presence of the most abundant strain, namely *Bartonella* sp. SC-tr1 (AJ583132). We found no significant effect of habitat type on the presence of *Bartonella* sp. SC-tr1 (df = 1, *p* = 0.83).

To evaluate the effect of habitat type on infection prevalence, we employed a GLM using the INLA framework. For *Bartonella*, we selected the model ‘*Bartonella ∼ Habitat + Sex + ELW’* that included both fixed and a spatial field (DIC = 65.15; Supplementary Table S4). The results indicated a significant effect of habitat type on *Bartonella* infection risk. Specifically, *Cricetomys* spp. captured in the urban environments showed markedly lower probabilities of infection (∼83%) compared to those from rural areas (posterior mean effect of urban habitat = −1.80, 95% CI: −3.12 to −0.48). This corresponds to a predicted infection probability of 26.5% (95% CI: 9.0–57.5%) in urban sites compared to 68.6% (95% CI: near 0.0–1.0%) in rural sites (Figure 4). Other covariates, i.e. sex and age, did not show a significant relation with infection status. Additionally, spatial structure was modelled using a stochastic partial differential equation (SPDE) approach, indicating no spatial autocorrelation in the *Bartonella* infection status (spatial variance component: σ^2^ = 0; Supplementary Figure S5). For *Hepatozoon*, we selected the model ‘*Hepatozoon ∼ Habitat + Sex + ELW’* that included both fixed effects and a spatial field (DIC = 45.73; Supplementary Table S4). No significant effect of habitat type on *Hepatozoon* infection prevalence was detected (posterior mean effect of urban habitat = −1.46, 95% CI: −3.20 to 0.27). However, our results showed a significant negative association between age and *Hepatozoon* infection prevalence (posterior mean effect of age (ELW) = −59.99, 95% CI: −98.41 to −21.56), implying that younger individuals are more likely to be infected. Notably, results of the spatial field analysis suggest possible autocorrelation in *Hepatozoon* infection (spatial variance component: σ^2^ = 0.3-0.36; Supplementary Figure S5), likely driven by the clustering of infected individuals from a single family group captured at the same location.

**Figure 4.**
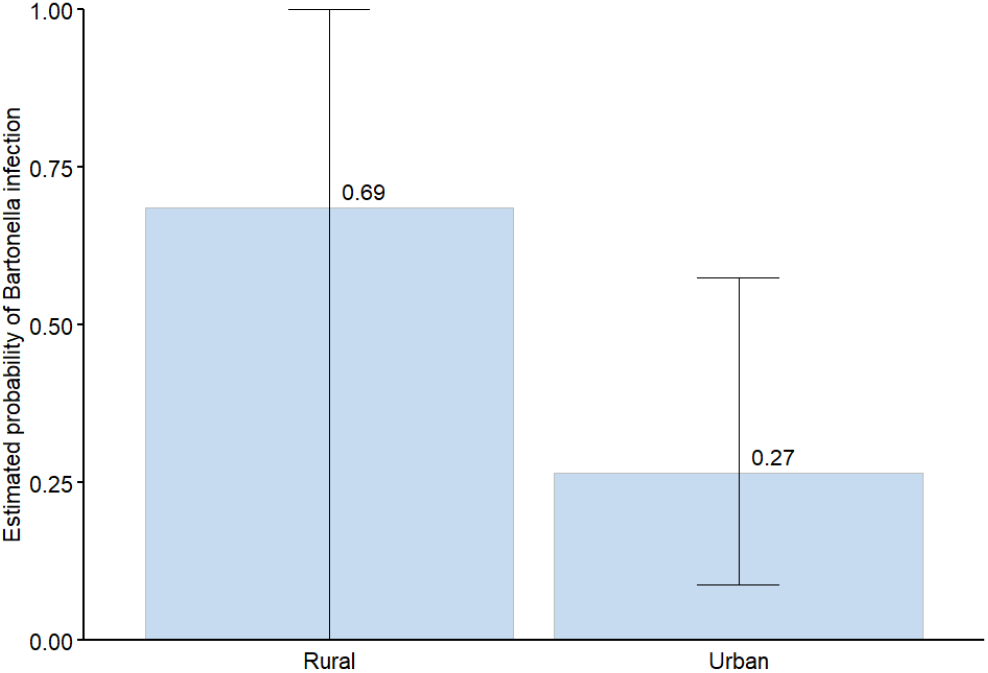
Estimated probability of *Bartonella* infection in *Cricetomys* spp. per habitat, presented with 95% confidence intervals.

Due to the limited number of positive samples, we did not conduct statistical analyses for *Anaplasma*.

## 4 Discussion

In this study, we investigated the prevalence of three microparasite genera in *Cricetomys* spp. from Morogoro, Tanzania, to assess the potential role of these rodents as reservoirs for zoonotic pathogens. We found a 71.9% prevalence of *Bartonella*, 17.5% for *Hepatozoon*, and 1.8% for *Anaplasma*.

We detected *Bartonella* spp. in 41 out of 57 *Cricetomys* spp. (71.9%, 95% CI: 60.2-83.6%), a higher prevalence than previously reported in sub-Saharan African rodents. For instance, a Tanzanian study detected an overall *Bartonella* prevalence of 17.0% in *Rattus rattus* using the *ssrA* gene and 5.0% using the *gltA* gene (Theonest et al. 2019). Another study reported a *Bartonella* prevalence of 61.0% in *Cricetomys gambianus* from Uganda using the *gltA* gene (Billeter et al. 2014). Additionally, previous research in Morogoro, Tanzania, reported a prevalence of 10.4% in *Mastomys natalensis* using the *gltA* and 16S-23S rRNA ITS region (Vanden Broecke et al. 2023). Of the 41 *Bartonella* sequences we obtained, 29 showed high similarity (97.5-99.3%) to sequences previously found in *M. natalensis* from the same research area (OQ506011, OQ506025, and OQ506048) (Vanden Broecke et al. 2023). This suggest that both *Cricetomys* spp. and *M. natalensis* may serve as reservoir hosts for the same or closely related *Bartonella* strains, potentially due to shared vectors. Furthermore, the phylogenetic analysis revealed a clustering of the obtained *Bartonella* sequences with sequences from Uganda and South Africa. This finding suggests that similar strains can infect different rodent hosts from diverse environments, highlighting the ecological flexibility and adaptability of *Bartonella*.

*Bartonella* infection prevalence is significantly influenced by habitat type, with the highest infection prevalence observed in rural areas compared to urban areas. Several factors may explain this outcome: (i) rural areas are characterized by a mosaic of agricultural land, fallow areas, and some human settlements, facilitating a higher degree of human-wildlife interaction and increasing the potential for reverse zoonosis from humans to animals. (ii) *Cricetomys* spp. captured in rural areas had a higher flea and tick load compared to individuals from urban areas. Given that fleas and ticks are known vectors for *Bartonella* spp. (Billeter et al. 2008), a higher parasitic load could result in higher infection prevalence. (iii) Urban habitats, where *Bartonella* infection prevalence was the lowest, may provide a more consistent food supply for rodents, allowing them to allocate more resources to their immune defences, thereby reducing susceptibility to infections (Budischak et al. 2017; Martin II et al. 2008). Nevertheless, it is important to note the wide 95% credible intervals associated with the estimated *Bartonella* infection probability in rural areas (0.0-1.0%), in contrast to the narrower intervals observed in urban areas (9.0–57.9%; Figure 4). This uncertainty likely reflects a combination of the small sample size and the ecological heterogeneity within the rural areas, encompassing transitional environments. Further research with larger sample sizes and a focus on microhabitat variability is necessary to fully understand the impact of habitat type on infection prevalence.

Despite these differences in infection prevalence, we found no significant variations in *Bartonella* strain compositions across the habitat types. This finding may be explained by the mobility of *Cricetomys* spp., which have a home range of approximately five hectares (Rey and Duplantier 2013). Their ability to move between different habitats may facilitate pathogen spillover, contributing to the homogenization of *Bartonella* diversity across different environments. While the spatial distribution and mobility of *Cricetomys* spp. are likely to influence pathogen transmission dynamics, we did not explicitly investigate spatial effects in this study.

*Hepatozoon* spp. were detected in 10 out of the 57 *Cricetomys* spp. (17.5%, 59% CI, 7.7-27.4%). This prevalence is higher than the 2.4% prevalence previously reported in *M. natalensis* in Tanzania (Vanden Broecke et al. 2023), but lower than the 41.0% prevalence found in rodents in North Africa (Maia et al. 2014). Additionally, all 10 *Hepatozoon* sequences exhibited high similarity to a sequence previously isolated from *M. natalensis* captured in the same research area (OL982745, 98.4-99.6%) (Vanden Broecke et al. 2023), suggesting that, similar to *Bartonella* strains, *Cricetomys* spp. and *M. natalensis* may serve as reservoir hosts for the same or closely related *Hepatozoon* strains. Furthermore, seven *Hepatozoon* sequences were most closely related to a *Hepatozoon* sp. isolated from the snake species *Pseudocerastes fieldi* in Iran (MZ412878) (Jameie et al. 2022), emphasizing that rodents can serve as intermediate hosts for snake hepatozoonosis (Dahmana et al. 2020). In fact, predation of rodents might be a crucial transmission pathway for *Hepatozoon* spp., as rodents consume invertebrates, and snakes can become infected by the ingestion of infectious rodents’ tissues (Sloboda et al. 2008).

Unlike *Bartonella*, we found no significant effect of habitat type on *Hepatozoon* infection prevalence. However, we observed a negative association between age and *Hepatozoon* infection status, implying higher infection rates among juveniles. This pattern may reflect age-related differences in immune responses, with younger individuals either more susceptible due to underdeveloped immunity or less able to clear infections effectively. Nonetheless, this relationship might be skewed because the small sample size of *Hepatozoon* positive individuals includes a high proportion of juveniles (6/10).

*Anaplasma* was found in one individual (1.8%, 95% CI, 0-5.2%). This prevalence is lower than the previously reported rate of 18.9% in *M. natalensis* in the same study area (Vanden Broecke et al. 2023). The identified sequence showed the highest similarity to an uncultured *Ehrlichia* sp. (OL982748, 98.8%) originally found in *M. natalensis* from the same research area (Vanden Broecke et al. 2023).

Using a spatially explicit Bayesian approach allowed us to investigate and control for spatial effects, addressing the spatial clustering of our sampling and the mobility of *Cricetomys* spp. Our analysis revealed no spatial autocorrelation in *Bartonella* infection, suggesting it may be widely distributed across the population, with limited influence from spatial covariates beyond our coarse habitat categorisation (urban vs. rural). However, the wide credible intervals in rural environments indicate greater uncertainty and highlight the need for further research in heterogeneous transitional landscapes (Figure 4). In contrast, Supplementary Figure S5 shows persistent spatial structuring for *Hepatozoon*, suggesting that its infection is localized in hotspots, potentially reflecting sample size and distribution effects.

One of the primary limitations of this study is the small sample size, necessitating cautious interpretation of the results. Additionally, we observed instances where multiple *Cricetomys* individuals were captured in the same cage during a single trapping night, often comprising familial groups. For instance, individuals CRI44, CRI45, and CRI46 were captured together. Both CRI45 and CRI46 tested positive for *Bartonella*. Similarly, individuals CRI50, CRI51, CRI52, CRI53, CRI54, CRI56, and CRI57 were captured together. Amongst these, apart from CRI52 and CRI56, all individuals tested positive for *Bartonella*, and all but CRI56 and CRI57 tested positive for *Hepatozoon*; this could therefore drive the spatial structuring we see in the results for *Hepatozoon* infections. Furthermore, individuals with uncertain infection status were excluded from all statistical analyses, potentially affecting the observed prevalences and distribution patterns.

## 5 Conclusion

This study investigated the prevalence of three microparasite genera in *Cricetomys* spp. in Morogoro, Tanzania. We found a 71.9% prevalence of *Bartonella*, 17.5% for *Hepatozoon*, and 1.8% for *Anaplasma*. These findings underscore the potential role of *Cricetomys* spp. as reservoirs for pathogens that may pose risks to humans, domestic animals, and livestock. Moreover, sequence analysis revealed a high similarity between *Bartonella, Hepatozoon*, and *Anaplasma* strains detected in *Cricetomys* spp. and those previously identified in *Mastomys natalensis* from the same research area, suggesting that both rodent species serve as reservoir hosts for the same or closely related strains. To our knowledge, this is the first study to investigate these microparasites in *Cricetomys* spp. in Tanzania.

Furthermore, our results provide preliminary evidence that *Bartonella* infection prevalence is influenced by habitat type, with significantly higher infection prevalence observed in rural areas compared to urban areas. However, further research with larger sample sizes is essential to fully understand the mechanisms driving these interactions and to develop effective strategies for managing and reducing the spread of infectious diseases from wildlife to humans and domestic animals.

## Supporting information

Supplementary table S1. Geographic coordinates of capture locations.

Supplementary figure S2. Phylogenetic tree of (A) Bartonella sequences, (B) Anaplasma and Ehrlichia sequences, and (C) Hepatozoon sequences.

Supplementary table S3. Sequence identity, host species, and geographical origin of Bartonella, Anaplasma, and Hepatozoon isolates.

Supplementary table S4. Results of the spatial GLM.

Supplementary figure S5. The spatial random effect predictions for Bartonella (top) and Hepatozoon (bottom) infection prevalence in Cricetomys spp.

## 7 Acknowledgement

We thank the staff (particularly Shabani Lutea, Geoffrey Sabuni, Omary Kibwana, and Ramadhani Idd Kigunguli) at the Institute of Pest Management (Sokoine University of Agriculture, Morogoro, Tanzania) for their excellent assistance during the fieldwork and dissections. We also thank two anonymous reviewers whose suggestions significantly improved the manuscript.

