## Supplementary table S1. Geographic coordinates of capture locations. for "Microparasite prevalence in Southern giant pouched rats (*Cricetomys ansorgei*) in Morogoro, Tanzania"

### Supplementary material

##### Supplementary table S1. Geographic coordinates of capture locations.

| Location | Habitat | Latitude | Longitude | Number of captured individuals |
|---|---|---|---|---|
| **House 1** | Urban | -6.811969 | 37.667170 | 7 |
| **House 2** | Urban | -6.822060 | 37.643130 | 3 |
| **Kitchen Garden** | Urban | -6.810970 | 37.666470 | 5 |
| **Garage** | Urban | -6.835550 | 37.665760 | 5 |
| **Kireka Bridge** | Rural | -6.855830 | 37.677780 | 4 |
| **Kireka Cornfield** | Rural | -6.861680 | 37.675730 | 2 |
| **Kireka School** | Rural | -6.865710 | 37.668870 | 2 |
| **Kireka Forest (edge)** | Forest | -6.888930 | 37.690740 | 4 |
| **Kireka Mosque** | Rural | -6.856000 | 37.674120 | 6 |
| **House 3** | Rural | -6.856370 | 37.675960 | 1 |
| **Kireka Bridge Upstream** | Rural | -6.855860 | 37.677727 | 2 |
| **Rock Garden** | Rural | -6.849863 | 37.673838 | 2 |
| **Kireka Village** | Rural | -6.855683 | 37.675530 | 2 |
| **House 4** | Urban | -6.814613 | 37.667878 | 1 |
| **Kiroka Mountain** | Rural | -6.858090 | 37.810010 | 11 |
