## Supplementary figure S2. Phylogenetic tree of (A) Bartonella sequences, (B) Anaplasma and Ehrlichia sequences, and (C) Hepatozoon sequences. for "Microparasite prevalence in Southern giant pouched rats (*Cricetomys ansorgei*) in Morogoro, Tanzania"

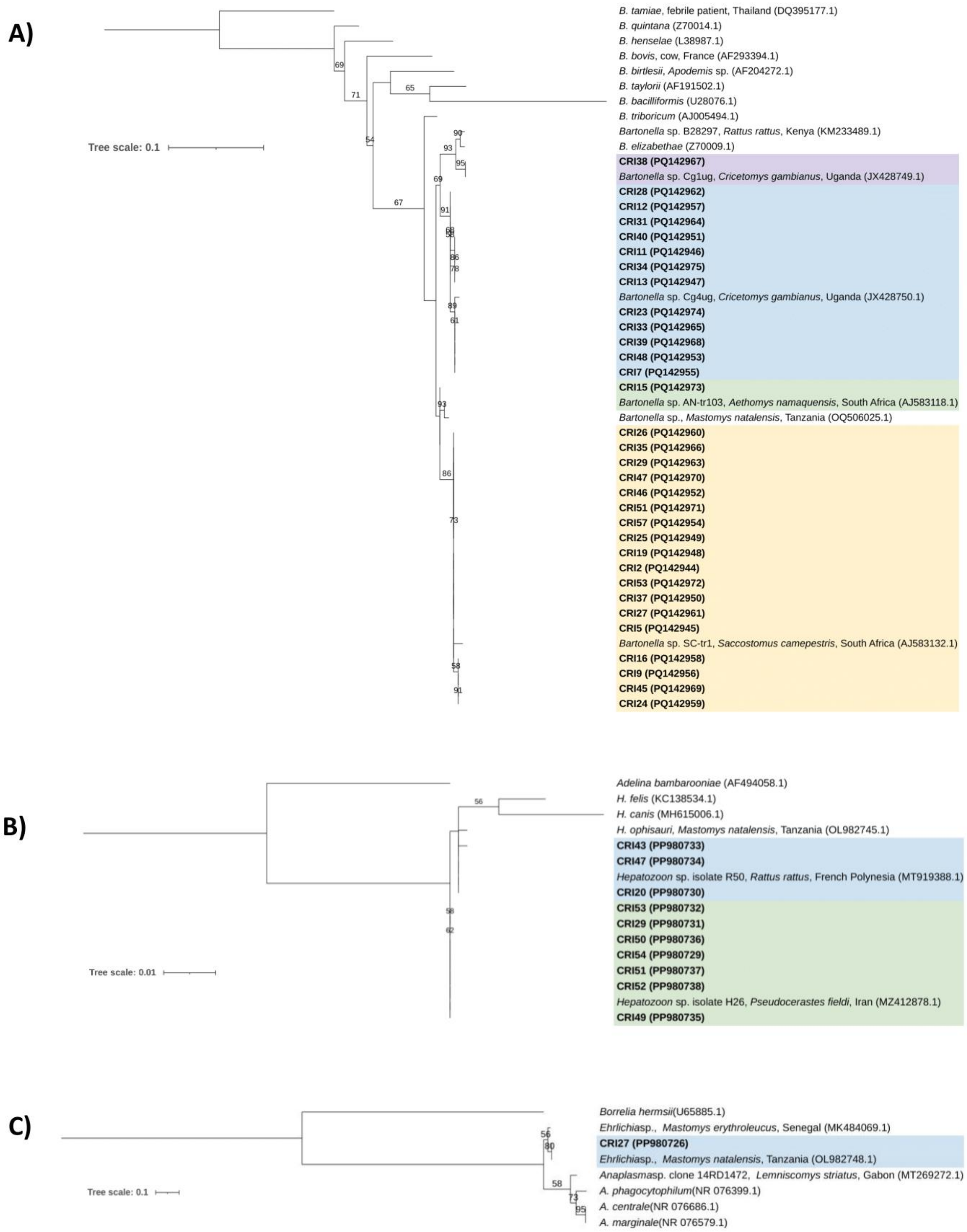


**Supplementary figure S2. Phylogenetic tree of (A) *Bartonella* gltA sequences, (B) *Anaplasma* and *Ehrlichia* 23S rRNA sequences, and (C) *Hepatozoon* 18S sequences derived from spleen tissue samples of *Cricetomys* spp. captured in Morogoro, Tanzania.** Bootstrap values greater than 50% are indicated above the corresponding branches. The sequences from this study are highlighted in bold and labelled with the prefix "CRI" followed by sample identifier numbers, and their respective GenBank accession numbers in parentheses. Reference strain sequences are included with the species name, host, and country, and GenBank accession numbers in parentheses. The colour of each sequence indicates the result of the BLAST analysis in GenBank, showing the highest similarity to the corresponding reference sequence. The outgroups used for the phylogenetic analysis were *B. tamiae* (DQ395177.1) for *Bartonella* spp., *Borrelia hermsii* for *Anaplasma* spp. (U65885.1), and *Adelina bambarooniae* (AF494058.1) for *Hepatozoon* spp.
