## Supplementary table S3. Sequence identity, host species, and geographical origin of Bartonella, Anaplasma, and Hepatozoon isolates. for "Microparasite prevalence in Southern giant pouched rats (*Cricetomys ansorgei*) in Morogoro, Tanzania"

**Supplementary table S3. Sequence identity, host species, and geographical origin of *Bartonella*, *Anaplasma*, and *Hepatozoon* isolates identified through Genbank BLAST analysis.** This table presents the results of BLAST hits for sequences obtained from *Cricetomys'* spleen samples, listing the corresponding GenBank accession numbers, organism, host species, geographical origin, and alignment statistics.

| Sample ID | First Hit (GenBank Accession number) | Organism | Host | Country | Grade | E Value | Length | % Identical Sites |
|---|---|---|---|---|---|---|---|---|
| **CRI2** | AJ583132 | *Bartonella* sp. | *Saccostomus campestris* | South Africa | 99.0% | 7.40E-135 | 280 | 98.2% |
| **CRI5** | AJ583132 | *Bartonella* sp. | *Saccostomus campestris* | South Africa | 99.2% | 9.17E-129 | 269 | 98.1% |
| **CRI7** | JX428750 | *Bartonella* sp. | *Cricetomys gambianus* | Uganda | 99.7% | 3.07E-123 | 251 | 99.2% |
| **CRI9** | AJ583132 | *Bartonella* sp. | *Saccostomus campestris* | South Africa | 99.2% | 3.70E-148 | 305 | 98.4% |
| **CRI10** | AJ583132 | *Bartonella* sp. | *Saccostomus campestris* | South Africa | 99.6% | 8.75E-56 | 128 | 98.4% |
| **CRI11** | JX428750 | *Bartonella* sp. | *Cricetomys gambianus* | Uganda | 99.2% | 6.15E-146 | 303 | 98.0% |
| **CRI12** | JX428750 | *Bartonella* sp. | *Cricetomys gambianus* | Uganda | 98.8% | 1.76E-110 | 237 | 97.9% |
| **CRI13** | JX428750 | *Bartonella* sp. | *Cricetomys gambianus* | Uganda | 99.4% | 3.77E-153 | 312 | 98.7% |
| **CRI15** | AJ583118 | *Bartonella* sp. | *Aethomys namaquensis* | South Africa | 99.1% | 3.86E-143 | 301 | 97.3% |
| **CRI16** | AJ583132 | *Bartonella* sp. | *Saccostomus campestris* | South Africa | 98.8% | 7.93E-145 | 299 | 98.3% |
| **CRI19** | AJ583132 | *Bartonella* sp. | *Saccostomus campestris* | South Africa | 98.9% | 3.84E-112 | 241 | 97.5% |
| **CRI20** | MT919388 | *Hepatozoon* sp. | *Rattus rattus* | French Polynesia | 99.7% | 0 | 564 | 99.5% |
| **CRI21** | HM636447 | *Bartonella massiliensis* | *Ornithodoros sonrai* | Senegal | 99.8% | 1.09E-70 | 153 | 99.3% |
| **CRI23** | JX428750 | *Bartonella* sp. | *Cricetomys gambianus* | Uganda | 99.7% | 7.57E-145 | 290 | 99.3% |
| **CRI24** | AJ583132 | *Bartonella* sp. | *Saccostomus campestris* | South Africa | 98.7% | 6.08E-141 | 296 | 97.6% |
| **CRI25** | AJ583132 | *Bartonella* sp. | *Saccostomus campestris* | South Africa | 99.1% | 1.71E-146 | 302 | 98.3% |
| **CRI26** | AJ583132 | *Bartonella* sp. | *Saccostomus campestris* | South Africa | 98.4% | 3.36E-154 | 337 | 95.5% |
| **CRI27** | OL982748 | Uncultured *Ehrlichia* | *Mastomys natalensis* | Tanzania | 99.9% | 0 | 406 | 99.8% |
| **CRI27** | AJ583132 | *Bartonella* sp. | *Saccostomus campestris* | South Africa | 99.3% | 8.72E-165 | 337 | 98.2% |
| **CRI28** | JX428750 | *Bartonella* sp. | *Cricetomys gambianus* | Uganda | 99.3% | 2.13E-140 | 291 | 97.9% |
| **CRI29** | AJ583132 | *Bartonella* sp. | *Saccostomus campestris* | South Africa | 99.2% | 6.76E-166 | 337 | 98.5% |
| **CRI29** | MZ412878 | *Hepatozoon* sp. | *Pseudocerastes fieldi* | Iran | 99.5% | 0 | 514 | 98.6% |
| **CRI31** | JX428750 | *Bartonella* sp. | *Cricetomys gambianus* | Uganda | 99.3% | 1.00E-143 | 296 | 98.3% |
| **CRI33** | JX428750 | *Bartonella* sp. | *Cricetomys gambianus* | Uganda | 99.8% | 1.96E-171 | 338 | 99.4% |
| **CRI34** | JX428750 | *Bartonella* sp. | *Cricetomys gambianus* | Uganda | 99.2% | 8.55E-119 | 250 | 98.4% |

| | | | | | | | | |
|---|---|---|---|---|---|---|---|---|
| **CRI35** | AJ583132 | *Bartonella* sp. | *Saccostomus campestris* | South Africa | 99.3% | 6.06E-146 | 299 | 98.7% |
| **CRI37** | AJ583132 | *Bartonella* sp. | *Saccostomus campestris* | South Africa | 99.3% | 3.36E-133 | 275 | 98.5% |
| **CRI38** | JX428749 | *Bartonella* sp. | *Cricetomys gambianus* | Uganda | 99.9% | 4.01E-173 | 338 | 99.7% |
| **CRI39** | JX428750 | *Bartonella* sp. | *Cricetomys gambianus* | Uganda | 99.7% | 2.41E-170 | 338 | 99.1% |
| **CRI40** | JX428750 | *Bartonella* sp. | *Cricetomys gambianus* | Uganda | 99.6% | 1.33E-152 | 307 | 99.0% |
| **CRI43** | MT919388 | *Hepatozoon* sp. | *Rattus rattus* | French Polynesia | 98.5% | 0 | 516 | 96.9% |
| **CRI45** | AJ583132 | *Bartonella* sp. | *Saccostomus campestris* | South Africa | 99.2% | 2.82E-144 | 299 | 98.3% |
| **CRI46** | AJ583132 | *Bartonella* sp. | *Saccostomus campestris* | South Africa | 99.3% | 4.88E-152 | 312 | 98.4% |
| **CRI47** | AJ583132 | *Bartonella* sp. | *Saccostomus campestris* | South Africa | 99.2% | 6.76E-166 | 337 | 98.5% |
| **CRI47** | MT919388 | *Hepatozoon* sp. | *Rattus rattus* | French Polynesia | 99.7% | 0 | 545 | 99.3% |
| **CRI48** | JX428750 | *Bartonella* sp. | *Cricetomys gambianus* | Uganda | 99.7% | 2.79E-149 | 298 | 99.3% |
| **CRI49** | MZ412878 | *Hepatozoon* sp. | *Pseudocerastes fieldi* | Iran | 100.0% | 0 | 548 | 100.0% |
| **CRI50** | AJ583132 | *Bartonella* sp. | *Saccostomus campestris* | South Africa | 98.6% | 1.56E-95 | 212 | 96.7% |
| **CRI50** | MZ412878 | *Hepatozoon* sp. | *Pseudocerastes fieldi* | Iran | 100.0% | 0 | 545 | 100.0% |
| **CRI51** | AJ583132 | *Bartonella* sp. | *Saccostomus campestris* | South Africa | 99.5% | 2.15E-145 | 295 | 99.0% |
| **CRI51** | MZ412878 | *Hepatozoon* sp. | *Pseudocerastes fieldi* | Iran | 99.96% | 0 | 559 | 99.8% |
| **CRI52** | MZ412878 | *Hepatozoon* sp. | *Pseudocerastes fieldi* | Iran | 100.0% | 0 | 512 | 100.0% |
| **CRI53** | AJ583132 | *Bartonella* sp. | *Saccostomus campestris* | South Africa | 99.0% | 4.25E-127 | 268 | 97.8% |
| **CRI53** | MZ412878 | *Hepatozoon* sp. | *Pseudocerastes fieldi* | Iran | 99.6% | 0 | 523 | 98.9% |
| **CRI54** | AJ583132 | *Bartonella* sp. | *Saccostomus campestris* | South Africa | 98.9% | 6.56E-115 | 247 | 97.6% |
| **CRI54** | MZ412878 | *Hepatozoon* sp. | *Pseudocerastes fieldi* | Iran | 100.0% | 0 | 534 | 100.0% |
| **CRI57** | AJ583132 | *Bartonella* sp. | *Saccostomus campestris* | South Africa | 99.2% | 2.83E-144 | 300 | 98.0% |
