## Supplementary table S4. Results of the spatial GLM. for "Microparasite prevalence in Southern giant pouched rats (*Cricetomys ansorgei*) in Morogoro, Tanzania"

**Supplementary table S4. Results of the spatial GLM analysing the effect of habitat type, sex, and ELW on the presence of *Bartonella* and *Hepatozoon* infection in *Cricetomys* spp.** This table presents the posterior means, standard deviation and lower (LCI) and upper (UCI) limits of the 95% credibility interval of the best-fitting INLA model as selected by minimising DIC. Terms are considered significant if the upper and lower credibility intervals do not cross zero (*).

| ***Bartonella*** **~ Habitat + Sex + ELW** | | | | |
|---|---|---|---|---|
| **Coefficients** | **Posterior mean** | **Standard deviation (SD)** | **Lower limit (LCI)** | **Upper limit (UCI)** |
| *Intercept* * | 0.781 | 23.879 | -46.654 | 47.938 |
| *HabitatUrban* * | -1.799 | 0.674 | -3.122 | -0.479 |
| *SexF* | -0.235 | 19.814 | -39.161 | 38.642 |
| *SexM* | 1.014 | 19.815 | -37.913 | 39.890 |
| *ELW* | -3.871 | 18.736 | -40.612 | 32.868 |
| DIC: 65.15 | | | | |
| ***Hepatozoon*** **~ Habitat + Sex + ELW** | | | | |
| **Coefficients** | **Posterior mean** | **Standard deviation (SD)** | **Lower limit (LCI)** | **Upper limit (UCI)** |
| *Intercept* | 0.449 | 26.172 | -50.879 | 51.775 |
| *HabitatUrban* | -1.463 | 0.885 | -3.198 | 0.272 |
| *SexF* | -0.022 | 20.529 | -40.278 | 40.233 |
| *SexM* | 0.402 | 20.528 | -39.853 | 40.655 |
| *ELW* * | -59.985 | 19.594 | -98.406 | -21.564 |
| AIC: 45.73 | | | | |
