## Supplementary figure S5. The spatial random effect predictions for Bartonella (top) and Hepatozoon (bottom) infection prevalence in Cricetomys spp. for "Microparasite prevalence in Southern giant pouched rats (*Cricetomys ansorgei*) in Morogoro, Tanzania"

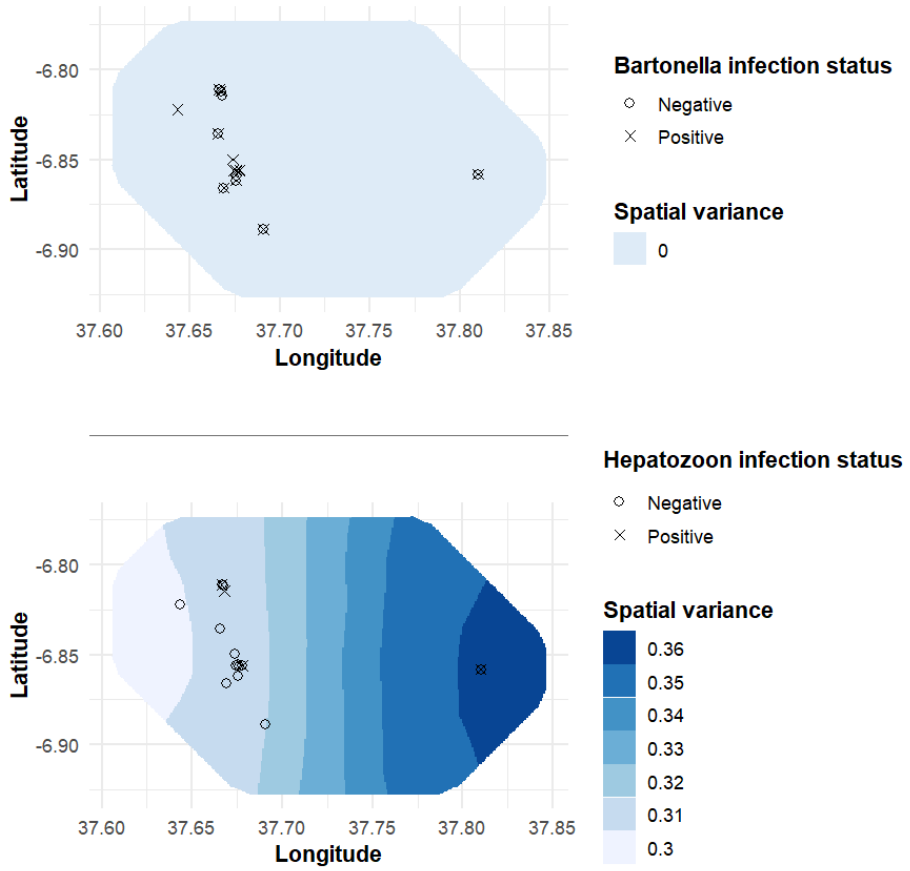


**Supplementary figure S5. The spatial random effect predictions for *Bartonella* (top) and *Hepatozoon* (bottom) infection prevalence in *Cricetomys* spp., generated using the SPDE-INLA framework.** The colour gradient represents the spatial field (spatial variance) across the study area, with darker shades indicating higher model-predicted variance. Points indicate individual rodent sampling locations, with symbols distinguishing infection status (positive versus negative).
